# GrgA as a potential target of selective antichlamydials

**DOI:** 10.1101/437285

**Authors:** Huirong Zhang, Sangeevan Vellappan, M. Matt Tang, Xiaofeng Bao, Huizhou Fan

**Affiliations:** Department of Pharmacology, Robert Wood Johnson Medical School, Rutgers University, Piscataway, New Jersey, USA; The George H. Cook Undergraduate Honors Scholars Program, School of Environmental and Biological Sciences, Rutgers University, New Brunswick, USA; Graduate Program in Physiology and Integrative Biology, Rutgers University, New Brunswick, USA; Department of Pharmacology, School of Pharmacy, Nantong University, Nantong, China

## Abstract

*Chlamydia* is a common pathogen that can causes serious complications in the reproductive system and eyes. Lack of vaccine and other effective prophylactic measures coupled with the largely asymptomatic nature and unrare clinical treatment failure calls for development of new antichlamydials, particularly selective antichlamydials without adverse effects on humans and the beneficial microbiota. We previously reported that benzal-N-acylhydrazones (BAH) can inhibit chlamydiae without detectable adverse effects on host cells and beneficial lactobacilli that dominate the human vaginal microbiota among reproductive-age women. However, the antichlamydial mechanism of BAH is not known. Whereas 4 single nucleotide polymorphisms (i.e., SNP1-4) were identified in a rare *Chlamydia* variant with a low level of BAH resistance, termed MCR, previous studies failed to establish a causal effect of any particular SNP(s). In the present work, we performed recombination to segregate the four SNPs. Susceptibility tests indicate that the R51G GrgA allele is both necessary and sufficient for the low level of BAH resistance. Thus, the *Chlamydia*-specific transcription factor GrgA either is a direct target of BAH or regulates BAH susceptibility. We further confirm an extremely low rate of BAH resistance in *Chlamydia*. Our findings warrant exploration of GrgA as a therapeutic and prophylactic target for chlamydial infections.

## INTRODUCTION

Chlamydiae are important and widespread pathogens. *Chlamydia trachomatis* is a leading infectious cause of blindness in many underdeveloped countries [1]. Globally, *C. trachomatis* is the leading sexually transmitted bacterial pathogen with an estimated prevalence of 4.2% among women aged 15-49 years [2]. In the United States, there has been a steep and sustained increase in sexually transmitted *C. trachomatis* infection in the past five years; 1.7 million cases were diagnosed in 2017, which represents a 22% increase from 2013, and accounts for 60% of cases of infections reported to the Centers for Disease Control and Prevention [3]. Genital *C. trachomatis* infection in women often leads to serious complications including infertility, pelvic inflammatory syndrome, abortion or premature birth and ectopic pregnancy [4].

*C. pneumoniae* is another common human pathogen, which causes bronchiolitis and pneumonia. Children, young adults and elderlies are at increased risks [5]. Several *Chlamydia* species are major health threats to livestock, and are also zoonotic pathogens [6, 7]. *C. muridarum* is a useful organism that models *C. trachomatis* infection in mice [8, 9].

Chlamydiae are susceptible to several broad-spectrum antibiotics. Human chlamydial infections are clinically treated with either azithromycin or doxycycline [10]. Due to a lack of vaccine, mass azithromycin administration has been used in Africa to treat eye infection and cut off the transmission. However, this chemical prevention strategy is only partially effective [11, 12]; furthermore, it has been linked to resistance development in standing-by pathogens [13, 14].

There are at least three additional concerns for current antichlamydial therapies. First, because of their broad-spectrums, they may cause dysbiosis in the genital tract and other systems [15–17]. Whereas loss of protective lactobacilli from the vagina of reproductive-age women may increase the risk of vaginal yeast infection [17], antibiotic-induced shift of gut microbiota may lead to problems ranging from severe diarrhea to increased risks for serious but not immediately noticeable metabolic changes [18, 19]. Second, although in culture *C. trachomatis* is highly susceptible to the therapeuticals, clinical treatment failure, which leads to persistent infection, is not rare [20, 21]. Finally, given the fact that tetracycline resistance has become widespread in *C. suis* due to farmers’ use of tetracycline as a growth promoter [22–24], antibiotic resistance could emerge in other *Chlamydia* species including *C. trachomatis* and *C. pneumoniae*.

For the above-mentioned reasons, it is important to identify new antichlamydial leads, particularly selective antichlamydial leads without adverse effects on either the host or other microbes, and identify their antichlamydial mechanisms. We have reported benzal-N-acylhydrazones (BAH) as novel antichlamydial leads capable of inhibiting all three *Chlamydia* species tested, *C. trachomatis, C. pneumoniae* and *C. muridarum* [25]. Significantly, at concentrations above minimal inhibition concentrations, BAH have no adverse effects on animal cells or vaginal lactobacilli [25]. Another attractive feature of BAH is their extremely low spontaneous mutation rates leading to resistance [25, 26]. Although a *C. muridarum* variant termed MCR with a low-level of BAH resistance was initially isolated following a lengthy selection process, multiple repeated attempts to isolate additional resistant variants from mutagenized as well as non-mutagenized stocks of *C. muridarum* and *C. trachomatis* were unsuccessful [25, 26].

How BAH inhibit chlamydiae remains unknown. Compared to the parental *C. muridarum*, MCR carries four single nucleotide polymorphisms (i.e., SNP1-4) in its genome (Table 1). SNP1 causes an A228V substitution in the major outer membrane protein (MOMP). Although A228 is conserved in MOMP in *C. muridarum* and *C. trachomatis*, V228 is found in *C. pneumoniae*, which remains highly susceptible to BAH [25]. SNP2 is located at the 10^th^ position of the 5’ untranslated region of the mRNA for Npt1 (ATP/ADP translocase), and is associated with a decreased Npt1 mRNA level. BAH have no effect on Npt1-mediated ATP transportation, suggesting that Ntp1 is unlikely a target of BAH [25]. SNP3 causes premature translation termination of TC0412, a homolog of the putative virulence factor CT135 in *C. trachomatis.* The truncated TC0412 contains only the N-terminal 23 amino acids, compared to the full length 365 amino acids. Given the hypermutable nature of *tc0412* [27] and the ultralow spontaneous BAH resistance rate, TC0412 is also unlikely a BAH target. Indeed, isogenic CT135 mutants are as susceptible to BAH as wild-type *C. trachomatis* [25]. SNP4 causes an R51G substitution in a *Chlamydia-*specific transcription activator termed GrgA. Whereas the transcription activation activity of GrgA is reduced by the substitution, it is not directly affected by BAH compounds [25]. Taken together, previous biochemical studies have failed to establish a role for MOMP, Npt1, TC0412 or GrgA in BAH-mediated *Chlamydia* inhibition. In this work, we establish through genome recombination that the rare R51G mutation in GrgA is both necessary and sufficient for BAH resistance in MCR. Our studies indicate GrgA as a promising target for selective antichlamydials.

**Table 1.** Terminology for SNPs in MCR.

|  | Effect | General term | Wild-type allele | Mutant allele |
| --- | --- | --- | --- | --- |
| SNP1 | A228V MOMP | S1(MOMP) | S1(wtMOMP) | S1(A228V MOMP) |
| SNP2 | Decreased Npt1 mRNA | S2(Npt1) | S2(wtNpt1) | S2(d_Npt1) |
| SNP3 | Truncated TC0412 | S3(TC0412) | S3(wtTC0412) | S3(t_TC0412) |
| SNP4 | R51G GrgA | S4(GrgA) | S4(wtGrgA) | S4(R51G GrgA) |

## Materials and Methods

### *Chlamydia* strains

Parental strains used for generation of recombinant chlamydiae as well as their precursors are listed in Table 2. Wild-type *C. muridarum* MoPn and the BAH-resistant variant MCR have been described previously [25]. MoPn_Rif^R^, MoPn_Spc^R^, MCR_LBM^R^ and MCR_Rif^R^ were derived by culturing MoPn and MCR in medium containing appropriate inhibitors (i.e., rifampin, spectinomycin or LBM415) at gradually increased concentrations starting at sub-MIC, as we previously outlined [25, 26, 28, 29].

**Table 2.**
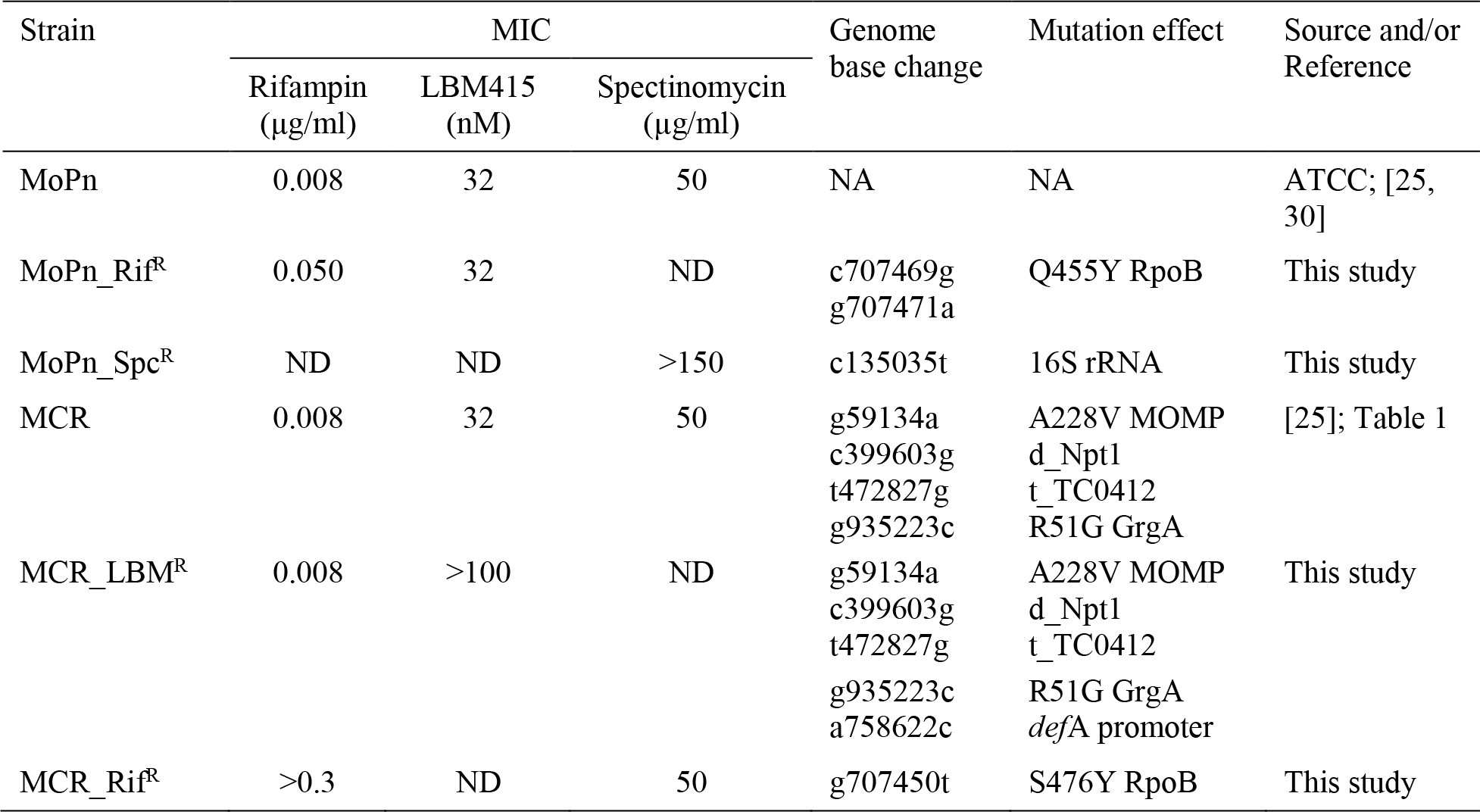
Strain information.

| Strain | MIC |  |  | Genome base change | Mutation effect | Source and/or Reference |
| --- | --- | --- | --- | --- | --- | --- |
|  | Rifampin (µg/ml) | LBM415 (nM) | Spectinomycin (µg/ml) |  |  |  |
| MoPn | 0.008 | 32 | 50 | NA | NA | ATCC; [25, 30] |
| MoPn_Rif<sup>R</sup> | 0.050 | 32 | ND | c707469g<br>g707471a | Q455Y RpoB | This study |
| MoPn_Spc<sup>R</sup> | ND | ND | >150 | c135035t | 16S rRNA | This study |
| MCR | 0.008 | 32 | 50 | g59134a<br>c399603g<br>t472827g<br>g935223c | A228V MOMP<br>d_Npt1<br>t_TC0412<br>R51G GrgA | [25]; Table 1 |
| MCR_LBM<sup>R</sup> | 0.008 | >100 | ND | g59134a<br>c399603g<br>t472827g<br>g935223c<br>a758622c | A228V MOMP<br>d_Npt1<br>t_TC0412<br>R51G GrgA<br>defA promoter | This study |
| MCR_Rif<sup>R</sup> | >0.3 | ND | 50 | g707450t | S476Y RpoB | This study |

Positions of SNPs in the genome are based on GenBank accession no. NC_002620. Abbreviation: MIC, minimal inhibitory concentration; NA, not applicable; Rif, rifampin; RpoB, RNA polymerase β subunit gene; Spc, spectinomycin; MOMP, major outer membrane protein; d_Npt1, decreased Npt1 (ATP/ADP translocase) expression; t_TC0412, truncated TC0412 protein with amino acids 1-23 (missing 24-365); *def*A, peptide deformylase gene; ND, not determined.

### Generation of recombinant chlamydiae

Mouse L929 cells grown in T25 flasks were coinfected with 2 parental strains at an MOI (multiplicity of infection) of 1 IFU per cell for each strain, cultured with medium containing 1μg/ml cycloheximide. After a passage without antibiotics, they were cultured with 6 ng/ml rifampin plus 25 nM LBM415 or 6 μg/ml spectinomycin (di selection) for 6 passages. 90 μM CF0001 was included as part of tri selection either following the completion of or in parallel to the di selection for 6 passages.

### Generation of clonal populations

Clonal populations of parental strains with resistance to rifampin, spectinomycin or LBM415 (Table 2) and recombinant chlamydiae were obtained mostly by limiting dilution [31] and in several cases by plaquing [32] following published protocols. When using limiting dilution, EB stocks were diluted to approximately 1 IFU per 96-well plate.

### Genotyping

Genomic DNA was prepared from infected cells using a Quick-gDNA™ MiniPrep kit (Zymo). DNA fragments for genes of interest were PCR-amplified, and sequenced at Genscript or MacrogenUSA using primers listed in Table S1 [25]. Peaks of sequencing chromatograms were manually checked for evidence for coexistence of wild-type and mutant alleles.

### Comparative BAH susceptibility tests

Near confluent HeLa cells were inoculated with chlamydiae at an MOI of 1 IFU per 10-30 cells, and cultured with medium containing 60 μM CF0001, indicated concentration of SF3, control solvent DMSO (final concentrations: 1.0% for CF0001 and 1.2% for SF3) and cycloheximide (1 μg/ml). 24 h later, cultures were harvested, and recoverable EB were quantified as previously described (22,23).

## RESULTS

### S4(R51G GrgA) is necessary for BAH resistance

To identify a particular SNP(s) that are necessary and/or sufficient for BAH resistance, we set out to segregate the 4 SNPs through genome recombination [33–35]. To enrich recombinant chlamydiae, we first derived a rifampin-resistant variant, termed MoPn_Rif^R^ from wild-type *C. muridarum* strain Nigg II (traditionally referred as strain mouse pneumonitis, MoPn), and an LBM415-resistant MCR variant, termed MCR_LBM^R^ (Table 2). Sequencing analyses revealed that the rifampin resistance in MoPn_Rif^R^ was due to two base changes in a single codon of the *rpo*B gene, resulting in an amino acid substitution (Q455Y) in the β subunit of the RNA polymerase (RpoB), whereas LBM415 resistance in MCR_LBM^R^ was due to a single point mutation in the promoter region of the *def*A (coding for peptide deformylase) [29], resulting in the generation of a predicted stronger −35 promoter element. As shown in Fig. S1, these mutations do not affect the antichlamydial effects of CF0001 [(*E*)-*N*’-(3,5-dibromo-4-hydroxybenzylidene)-3-dinitrobenzohydrazide], a prototype BAH.

We performed two MoPn_Rif^R^ X MCR_LBM^R^ recombination studies. For the first one, we coinfected 5 flasks of L929 cells with the two parental strains, and maintained the flasks as independent lines (W1-5) in subsequent passages (Fig. 1A). We selected for recombinant chlamydiae using sub-minimal inhibitory concentrations of rifampin and LBM415 (see experimental procedures). At the end of the 6^th^ passage of the Rif/LBM di selection, Sanger’s sequencing revealed that wild-type *rpo*B and *def*A alleles were apparently eliminated in 4 of the 5 lines (Fig. 1B), whereas the W4 line still retained both the wild-type and mutant alleles of *rpo*B and *def*A. These results indicate that at this point the W1, W2, W3 and W5 lines were comprised of recombinants and very few (if any) parental organisms (Fig. 1B). Contrast to the *rpo*B and *def*A selection markers, almost all loci of the 4 SNPs displayed a mixture of wild-type and mutant alleles in these lines (Fig. 1B), suggestive of good recombination complexity at most of the SNP loci. Exceptions were apparent absence of S1(A228V MOMP) and S2(wtNpt1) alleles in the W1 and W3 lines, respectively (Fig. 1B), likely reflecting low recombination complexity at these sites in these lines. We then continued the selection for BAH resistance by adding CF0001 to the Rif/LBM di selection for 6 additional passages. Interestingly, by the end of the 6^th^ passage with the Rif/LBM/CF tri selection, we observed apparent elimination of wild-type alleles at all the 4 SNP loci in all 5 lines (Fig. 1C), even for the locus of S1(MOMP), where the S1(A228V MOMP) allele was unnoticeable (and thus must be present at a very low percentage) prior to the start of the tri selection (Fig. 1B).

**Fig. 1.**
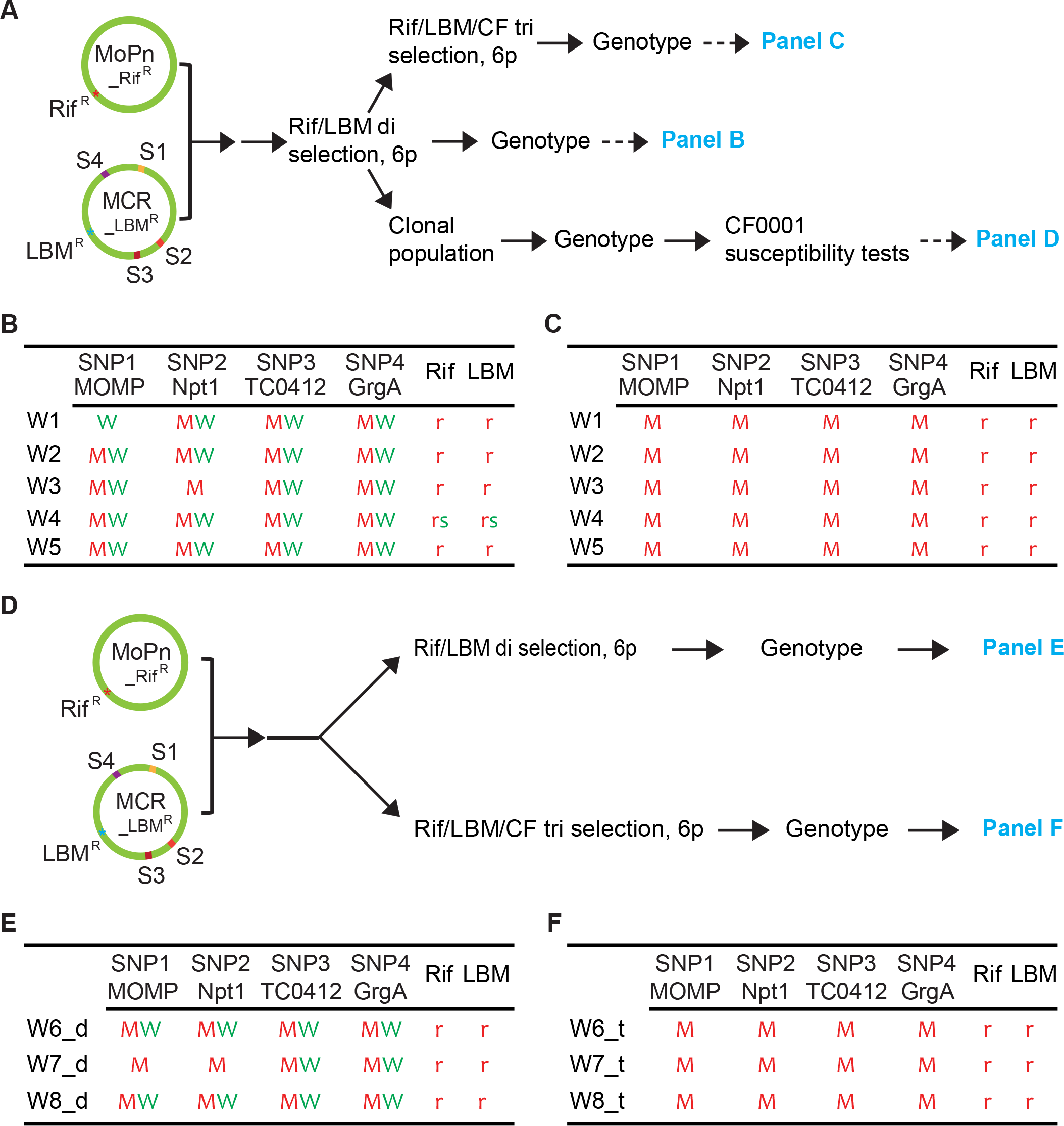
The Rif/LBM di selection is largely non-discriminatory towards either allele at the SNP loci, but the Rif/LBM/CF tri selection eliminates wild-type alleles. (A) Schematic presentation of genomes of MoPn_Rif^R^ (*C. muridarum* MoPn variant resistant to rifampin) and MCR_LBM^R^ (derivative of MoPn variant MCR with resistance to LBM415) and experimental flow for recombination, sequential di and tri selection, genotyping, susceptibility tests and data presentation. (B) Genotyping results of Rif/LBM-selected populations from the first MoPn_Rif^R^ X MCR_LBM^R^ recombination study. (C) Genotyping results of Rif/LBM/CF-selected populations showing elimination of wild-type alleles at all SNP loci by CF0001 from Rif/LBM-selected populations in (B). (D) Experimental flow for MoPn_Rif^R^ X MCR_LBM^R^ recombination, parallel di and tri selection, genotyping, susceptibility tests and data presentation. (E) Genotyping results of Rif/LBM-selected populations of the second MoPn_Rif^R^ X MCR_LBM^R^ recombination study. (F) Genotyping results of Rif/LBM/CF-selected populations obtained in parallel to those in (E). (A, D) S1, S2, S3 and S4 signify the four SNPs. Rif, rifampin; LBM, LBM415; CF, CF0001; 6p, 6 passages. (B, C, E, F) Green “W” and red “M” signify a wild-type allele and a mutant allele, respectively, at the SNP loci. Green “s” and red “r” signify wild-type and mutated genotypes, which render susceptibility and resistance, respectively, to either rifampin or LBM415.

For the second recombination study, we co-infected 3 flasks of L929 cells with MoPn_Rif^R^ and MCR_LBM^R^. We modified the selection protocols by splitting each line into two fractions, and subjecting them to the Rif/LBM di selection and Rif/LBM/CF tri selection in parallel (as opposed to sequential di and tri selections above) (Fig. 1D). By the end of 6^th^ passages with either selection regimen wild-type *rpo*B and *def*A alleles were apparently absent, indicating (near) complete elimination of parental chlamydiae (Fig. 1E). In Rif/LBM-selected populations, both wild-type and mutant alleles were present at most of the SNP loci (Fig. 1E). In comparison, Rif/LBM/CF-selected populations exhibited only mutant alleles at all the loci (Fig. 1F). The consistent elimination of chlamydiae carrying any wild-type alleles by CF0001 in both recombination studies suggests two alternative probabilities. In one, all of the 4 mutant alleles in MCR are necessary for BAH resistance. In the other, only 1 or up to 3 of the 4 mutant alleles are necessary, but accompanying mutant allele(s) compensate for growth disadvantages caused by the mutant allele(s) required for BAH resistance.

To distinguish between these probabilities, we set out to generate clonal populations from the W1, W2, W3 and W5 lines that were subjected to 6 passages of Rif/LBM di selection through either limiting dilution [31] or plaquing [32]. A total of 79 clonal populations were generated. Complete genotyping analyses were performed for 32 populations, which represented only 8 of the 16 possible recombinant genotypes (Table S2). The remaining 47 clonal populations were only partially genotyped because initial sequencing data indicated that either they were likely redundant populations or their genotypes were considered unhelpful based on BAH susceptibility data that were already obtained from fully genotyped populations.

BAH susceptibility tests were performed for 14 clonal populations, which represented all of the 8 defined unique genotypes, alongside with MCR and MoPn (Fig. 2). As expected, both all-wild-type allele populations tested (i.e., w5c2 and w1c15) displayed susceptibility to CF0001 as wild-type MoPn, whereas both all-mutant-allele populations tested (i.e., w3c2 and w5c4) displayed resistance as MCR. Interestingly, among 10 clonal populations with 1-3 mutant alleles, only w2c10, the sole clonal population with an S4(R51G GrgA) allele, was resistant, whereas all 9 other populations, which carried S4(wtGrgA) were susceptible even though they had up to 3 mutant alleles at S1(MOMP), S2(Npt1) and/or S3(TC0412). These results suggest that among the 4 SNPs in MCR, only S4(R51 GrgA) is required for BAH resistance. However, due to the coexistence of the S3(t_TC0412) allele in w2c10, and the lack of a population with S4(R51G GrgA) as the only mutant allele, it was unclear whether the R51G GrgA allele alone is sufficient for BAH resistance.

**Fig. 2.**
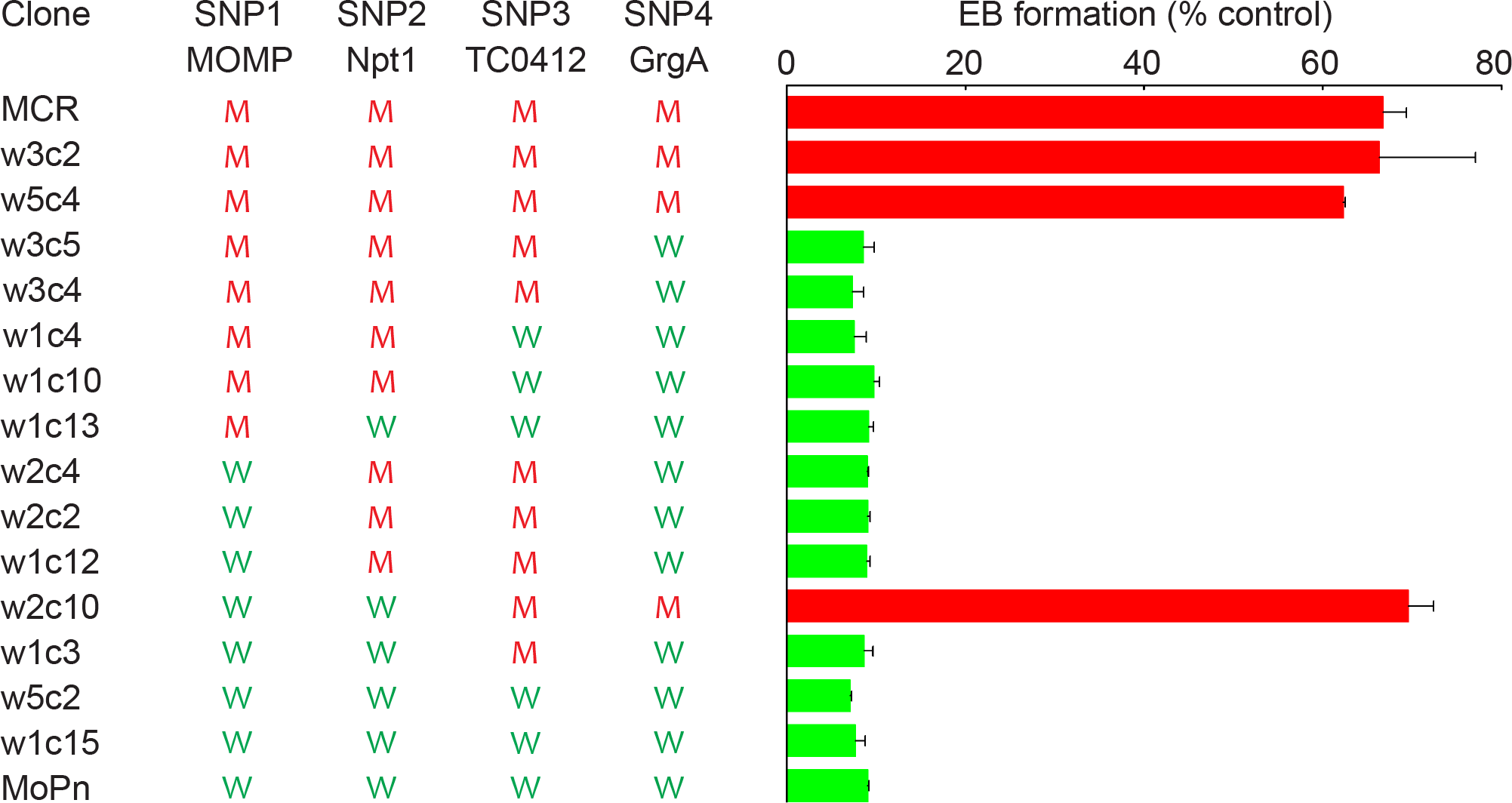
S4(R51G GrgA) is necessary for BAH resistance. CF0001 inhibition profile of representative fully genotyped clonal populations. Green “W” and red “M” signify a wild-type allele and a mutant allele, respectively. EB formation was determined in medium containing either 60 μM CF0001 or 1% DMSO as vehicle control. Results were averages ± standard deviation of triplicate experiments.

### SNP4(R51G GrgA) is sufficient for BAH resistance

In the above MoPn_Rif^R^ X MCR_LBM^R^ recombination studies, the selection markers *rpo*B and *def*A are both located between S3(TC0412) and S4(GrgA) in the MoPn genome (Fig. 1A, D). To obtain variants with a genotype of S1(wtMOMP), S2(wtNpt1), S3(wtTC0412) and S4(R51G GrgA), we decided to use two selection markers separated by a SNP(s). We derived a spectinomycin-resistant *C. muridarum* variant (MoPn_Spc^R^), which carries a single point mutation in the 16S rRNA (Table 3). Unlike *rpo*B and *def*A, the 16S rRNA gene is located between S1(MOMP) and S2(Npt1) (Fig. 3A). This mutation did not affect inhibition of chlamydiae by CF0001 (Fig. S1).

**Fig. 3.**
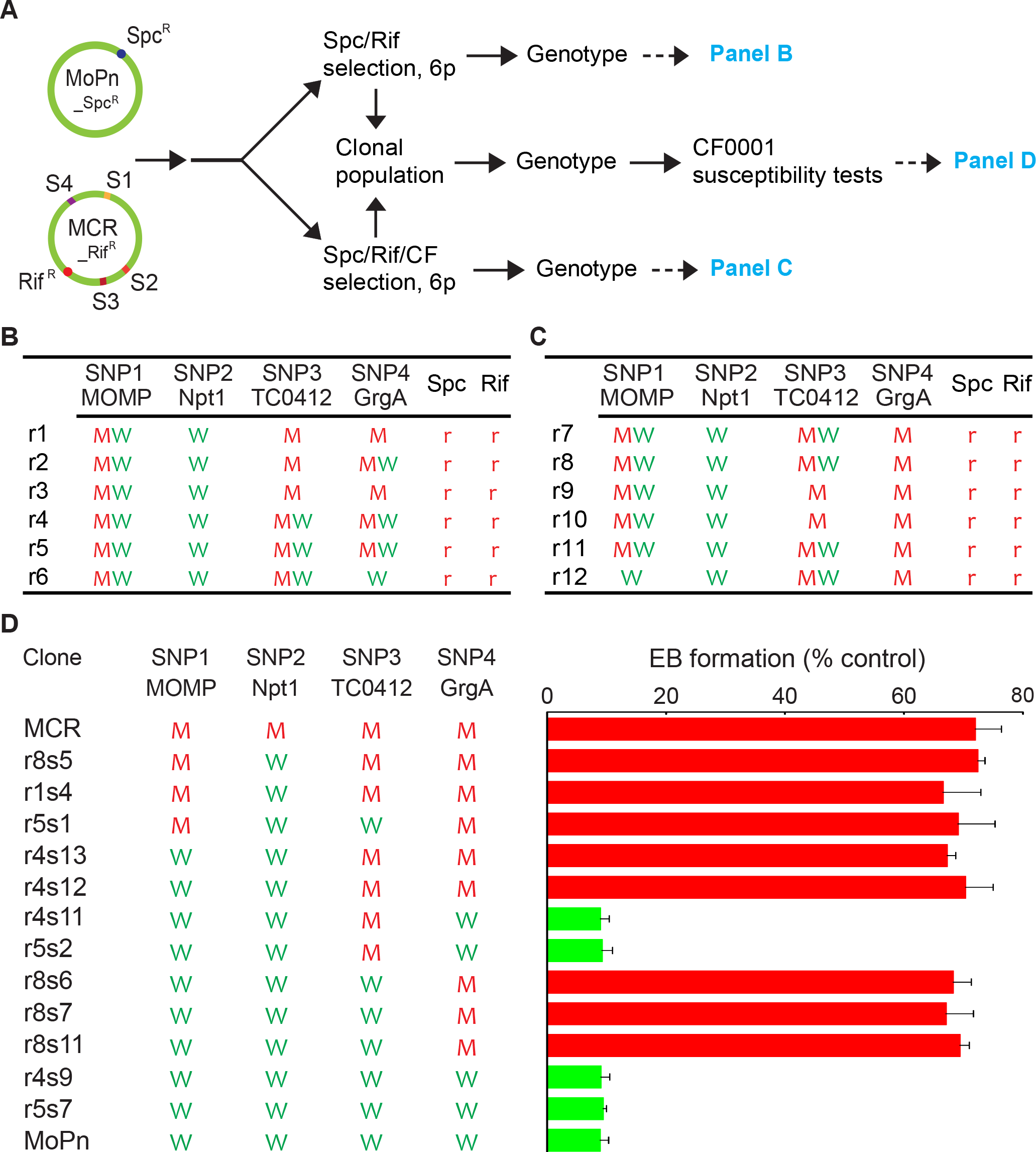
S4(R51G GrgA) is sufficient for BAH resistance. (A) Schematic presentation of genomes of MoPn_Spc^R^ (*C. muridarum* MoPn variant resistant to spectinomycin) and MCR_Rif^R^ (derivative of MoPn variant MCR with resistance to rifampin) and experimental flow for recombination, recombinant selection, genotyping, susceptibility tests and data presentation. (B) Genotyping results of Spc/Rif-selected populations. (C) Genotyping results of Spc/Rif/CF-selected populations. (D) CF0001 inhibition profiles of representative fully genotyped clonal populations as determined in Fig. 2. (A-D) Refer to Fig. 1 legend for signification and abbreviations.

Initially, we tried to generate but failed to select for MoPn_Spc^R^ X MCR_LBM^R^ recombinants using the Spc/LBM (spectinomycin plus LBM415) selection regimen in 3 different attempts. In the regimen, the concentration of LBM415 was either the same as or higher than the concentration used for the above Rif/LBM regimen. The Spc/LBM regimen failed to eliminate wild-type *def*A allele although it successfully eliminated wild-type 16S rRNA. These findings suggest that spectinomycin and LBM415 are incompatible for recombinant selection.

We next derived an MCR variant with rifampin resistance (MCR_Rif^R^). Similar to MoPn_Rif^R^, which expresses a Q455Y RpoB, MCR_Rif^R^ expresses an S476Y RpoB (Table 3). MCR_Rif^R^ retained low level of CF0001-resistance as MCR and MCR_LBM415 (Fig. S1).

We created 6 independent MoPn_Spc^R^ X MCR_Rif^R^ recombinant lines. Each of the 6 lines was subjected to parallel Spc/Rif di selection and Spc/Rif/CF tri selection, and subsequently to genotyping analyses (Fig. 3A). After 6 passages with either selection, most (if not all) parental chlamydiae were eliminated, as indicated by apparent lack of the spectinomycin-susceptible 16S rRNA allele and rifampin-susceptible *rpo*B allele (Fig. 3B, C).

Genotyping analyses suggest that allele diversity for the 4 SNPs in the Spc/Rif-selected MoPn_Spc^R^ X MCR_Rif^R^ recombinants (Fig. 3B) was lower than that of the Rif/LBM-selected MoPn_Rif^R^ X MCR_LBM^R^ recombinants (Fig. 1B). Although the Spc/Rif di selection displayed no bias for either S1(MOMP) alleles, it showed a consistent bias for the S2(wtNpt1) in all 6 lines. It also displayed bias for the S3(t_TC0412) alleles in 3 (r1-3) of the 6 lines. For the S4(GrgA) locus, it displayed bias for (in lines r1 and r3) and against (in line r6) the S4(R51G GrgA) allele, and no apparent bias in the remaining 3 lines (r2, r4 and r5).

In a striking contrast to the Rif/LBM/CF-selected MoPn_Rif^R^ X MCR_LBM^R^ recombinants, which consistently carried only mutant alleles at all 4 SNP loci (Fig. 1C), Spc/Rif/CF-selected MoPn_Spc^R^ X MCR_Rif^R^ recombinants contained both S1(wtMOMP) and/or S3(wtTC0412) alleles, in addition to mutant alleles, at the S1(MOMP) and S3(TC0412) loci, but only wild-type allele at the S2(Npt1) locus (Fig. 3C). The only consistency between Rif/LBM/CF-selected populations and Spc/Rif/CF-selected populations is the lack of wild-type allele at the S4(GrgA) locus (Fig. 3C), further supporting the notion that the S4(R51G GrgA) allele is necessary for BAH resistance (Fig. 2).

We generated 21 clonal populations from Spc/Rif-selected populations, and 13 clonal populations from Spc/Rif/CF-selected populations. These 34 clonal populations represented 6 of the 16 possible recombinant genotypes (Table S3). 12 representative clonal populations were tested for BAH resistance alongside with MCR and MoPn. All 4 clonal populations carrying the S4(wtGrgA) allele demonstrated susceptibility to CF0001, whereas all 8 clonal populations carrying the S4(R51G GrgA) allele including the 3 populations (r8s6, r8s7 and r8s11) carrying S4(R51G GrgA) and wild-type alleles for the 3 remaining SNP loci were resistant (Fig. 3D). These findings, together with data presented in Fig. 2, indicate that the S4(R51G GrgA) allele is both necessary and sufficient for BAH resistance, which can be viewed more clearly by arranging all phenotypically characterized clonal populations from both the MoPn_Rif^R^ X MCR_LBM^R^ recombination and the MoPn_Spc^R^ X MCR_Rif^R^ recombination by their S4(GrgA) genotype (Table S4).

### Ultralow rate of spontaneous BAH resistance

Previous studies indicated that BAH resistance in chlamydiae occurs at extremely low rates. The observations that the Rif/LBM/CF selection consistently eliminated wild-type S1(wtMOMP), S2(wtNpt1) and S3(wtTC0412) (Fig. 1C, F) even though S4(R51G GrgA) is the only mutant allele that is required for BAH resistance suggest that accompanying mutations help the survival of chlamydiae carrying S4(R51G GrgA) in the presence of BAH on the background of mutated *rpo*B and *def*A. We next determined whether presence of mutant alleles at the S1(MOMP), S2(Npt1) and S3(TC0412) loci in the genome helps selection for variants with GrgA mutation conferring BAH resistance by using the clonal population w3c5, which carries wild-type GrgA allele at the S4(GrgA) locus but mutant alleles at all three remaining SNP loci.

A total of 6 screens were carried out with w3c5. The first 2 screens were initiated with a combined 0.9 × 10^7^ inclusion-forming units (IFU) of non-mutagenized elementary bodies (EB) and selection was carried out with CF0001 (gradually increased from 80-120 μM) as a sole selection agent. The second 2 screens were initiated with the same number of non-mutagenized EB but selection was carried out with the Rif/LBM/CF tri selection regimen that was used to select for CF0001-resistant recombinants (Fig. 1A, D). No resistant chlamydiae emerged with either selection regimen.

The final two screens for CF0001-resistant variants were initiated with EBs prepared from cultures treated with 2 or 5 mg/ml ethyl methanesulfonate, a DNA-damaging reagent that has been used to mutagenize chlamydiae previously [29, 36, 37]. We also failed to obtain resistant chlamydiae in each of these attempts starting with 2 × 10^7^ IFUs of EB. The repeated failures to isolate additional CF0001-resistant variants suggest that only very few and specific mutations in GrgA can lead to BAH resistance and/or sustain chlamydiae.

### S4(R51G GrgA)-mediated BAH resistance is overcome by SF3

Compared to the prototype antichlamydial BAH CF0001, SF3 [(*E*)-*N*’-(3,5-dibromo-4-hydroxybenzylidene)-3,5-dinitrobenzohydrazide], a recently developed BAH, has a stronger antichlamydial activity, and can fully inhibit MCR [26]. At 80 μM, SF3 also achieved full inhibition of all three tested clonal populations (r8s6, r8s7 and r8s11) with S4(R51G GrgA) as the sole mutant allele although at lower concentrations it inhibited the wild-type MoPn more efficiently (Fig. 4). These results indicate that S4(R51G GrgA)-mediated BAH resistance can be overcome by SF3.

**Fig. 4.**
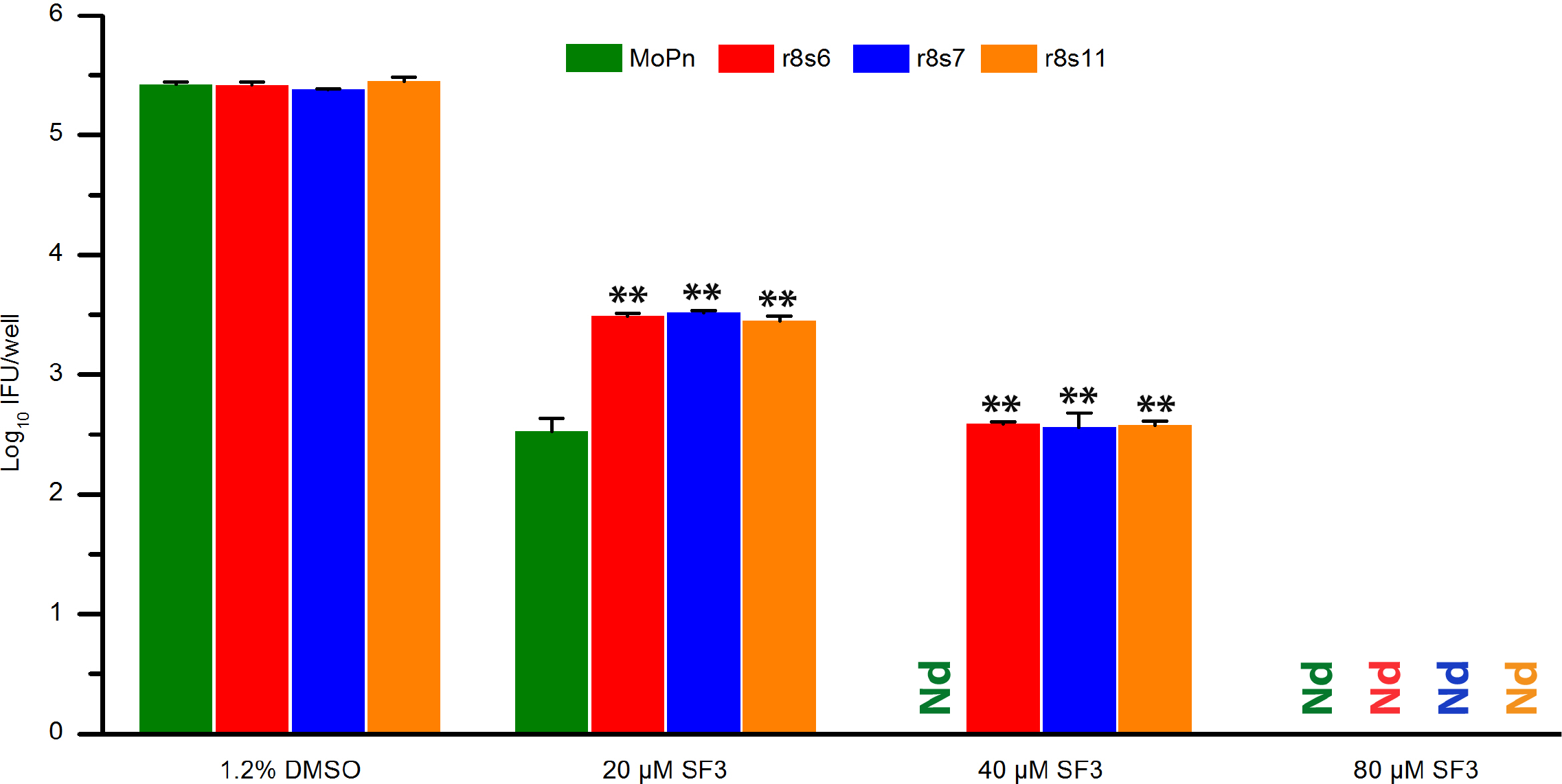
Complete inhibition of clonal populations with S4(R51G GrgA) as the sole mutant allele by 80 μM SF3. Results are averages ± standard deviations of duplicate experiments. Nd, none detected. Double asterisks signify statistically significantly higher number of EBs formed by the clonal recombinant populations, as compared to wild-type MoPn, in the presence of 20 or 40 μM SF3 (*p* < 0.01, 2-tailed *t* test).

## DISCUSSION

BAH belong to a novel group of selective antichlamydials [25, 26]. The genome of the rare BAH-resistant *C. muridarum* variant MCR carries four SNPs [25]. Through extensive genetic analyses and susceptibility tests reported here, we have unequivocally established that the S4(R51G GrgA) allele is both necessary and sufficient for a low level of BAH resistance.

GrgA functions as a transcription activator. Although it has been previously demonstrated that BAH is incapable of blocking the transcription activation activity of GrgA *in vitro* [25], it is possible that BAH functions as “prodrug”; host cell-or chlamydia-derived BAH derivatives may interact with GrgA. Alternatively, BAH may interfere with another yet-to-be defined critical process involving GrgA. It is also conceivable that GrgA regulates BAH susceptibility without directly interacting with BAH or their bioactive derivatives.

*Chlamydia* encodes 3 sigma factors, including the major sigma factor σ^66^ and two alternative sigma factors σ^28^ and σ^54^. As part of the RNA polymerase, the sigma factors recognize different promoter sequences. Studies have shown that GrgA activates both σ^66^-dependent transcription and σ^28^-dependent transcription *in vitro*, suggestive of critical roles for GrgA in chlamydial gene expression [38, 39]. Thus, GrgA is a promising candidate therapeutic and prophylactic target even if it may not be the receptor of BAH or their bioactive derivatives. GrgA is a *Chlamydia-* specific protein. Whereas it is conserved by all *Chlamydia* species, it is not found in any other organisms. Therefore, targeting GrgA will provide intrinsically high selectivity.

Previous studies have shown that random mutation rates leading to BAH resistance is extremely low in *C. trachomatis* and *C. muridarum*. Therefore, MCR represents a rare variant with only a low level of resistance. Consistent elimination of wild-type alleles at the loci of SNP1-3 from Rif/LBM/CF-selected populations suggests that the co-existence of these mutant alleles helps survival of chlamydiae carrying the S4(R51G GrgA) allele in the presence of BAH, rifampin and LBM415. Our failures to isolate additional BAH-resistant mutants on the background of S1(A228V MOMP), S2(d_Npt1) and S3(t_TC0412) from clonal population w3c5 in multiple attempts with different selection regimen suggest that very few and specific GrgA mutations can cause BAH resistance and/or sustain chlamydial growth.

Compared to the prototype antichlamydial BAH CF0001, the recently-developed SF3 has a stronger antichlamydial activity while maintaining non-toxicity to mammalian cells and vaginal lactobacilli [26]. It has been shown previously that MCR can be inhibited completely by SF3 even though it is less susceptible than wild-type MoPn to lower concentrations of SF3 [26]. While it is expected that clonal recombinant populations carrying S4(R51G GrgA) as sole mutant allele demonstrate the same properties as MCR, these clonal populations will be more useful for identifying additional selective antichlamydials that interferes with a process involving GrgA.

In summary, we have unequivocally established that R51G GrgA is both necessary and sufficient for the low level of BAH resistance in the *Chlamydia* variant MCR. These findings and the facts that GrgA is a *Chlamydia-*specific protein and plays important roles in chlamydial transcription indicate GrgA as a promising selective therapeutic/prophylactic target, even though it is unclear whether GrgA is a direct target of BAH or regulates BAH susceptibility without directly interacting with BAH. In addition to the high selectivity, the ultralow rate of BAH resistance in chlamydiae is another super attractive feature for developing BAH compounds as therapeutic/prophylactic agents.

## ACKNOWLEDGEMENTS

We thank Prof. Daniel Seidel (University of Florida) for supplying CF0001 and SF3, Novartis Institutes for BioMedical Research for supplying LBM415, and Prof. Spencer Knapp (Rutgers University) for helpful discussions. This work was supported by National Institutes of Health (Grant # AI122034 to HF), New Jersey Health Foundation (Grant # PC 20-18 to HF), Natural Sciences Foundation of China (grant # 31400165 to XB). SV was a Rutgers Aresty Research Scholar in 2015 and 2016.

